# White Matter Brain Structure Predicts Language Performance and Learning Success

**DOI:** 10.1101/2022.01.14.476338

**Authors:** Stella M. Sánchez, Helmut Schmidt, Guillermo Gallardo, Alfred Anwander, Jens Brauer, Angela D. Friederici, Thomas R. Knösche

## Abstract

Individual differences in the ability to deal with language have long been discussed. The neural basis of these, however, is yet unknown. Here we investigated the relationship between long-range white matter connectivity of the brain, as revealed by diffusion tractography, and the ability to process syntactically complex sentences in the participants’ native language as well as the improvement thereof by multi-day training. We identified specific network motifs that indeed related white matter tractography to individual language processing performance. First, for two such motifs, one in the left and one in the right hemisphere, their individual prevalence significantly predicted the individual language performance suggesting a predisposition for the individual ability to process syntactically complex sentences, which manifests itself in the white matter brain structure. Both motifs comprise a number of cortical regions, but seem to be dominated by areas known for the involvement in working memory rather than the classical language network itself. Second, we identified another left hemispheric network motif, whose change of prevalence over the training period significantly correlated with the individual change in performance, thus reflecting training induced white matter plasticity. This motif comprises diverse cortical areas including regions known for their involvement in language processing, working memory and motor functions. The present findings suggest that individual differences in language processing and learning can be explained, in part, by individual differences in the brain’s white matter structure. Brain structure may be a crucial factor to be considered when discussing variations in human cognitive performance, more generally.

## 1. Introduction

Language is a cognitive domain that is considered specifically human, in particular when it comes to processing syntactically complex structures (Berwick and Chomsky, 2016; Fitch and Hauser, 2004; Hauser et al., 2002). People, however, differ in how well they deal with processing complex sentences, even in their native language, depending on their working memory capacity (Caplan and Waters, 1999; MacDonald et al., 1992). When this capacity is low, processing of ambiguous sentences becomes difficult (Fiebach et al., 2004; Friederici et al., 1998; Just and Carpenter, 1992). Independently, it has been shown that one’s capability to process complex sentences in the native language can be improved in relatively short time by intense training, even in adults (Wang et al., 2021). This raises two questions: What is the neurobiological basis of behavioral differences in processing complex sentences, and what is the neural basis responsible for training-induced performance improvements in processing such sentences.

The neuroanatomical and physiological underpinnings of language processing and language learning are certainly diverse. Language processing in the adult brain is mainly based on a left hemispheric network involving particular frontal, temporal and parietal regions (for a review see Friederici, 2011). By contrast, language learning – at least second language learning in adults – appears to involve additional brain regions (for a review see Li et al., 2014). The multiple spatially separated regions involved in language processing are connected by dorsally and ventrally located long-range fiber bundles running through the white matter of the brain (for an overview, see Friederici, 2017). Among these, the dorsal fiber tract targeting Broca’s area is particularly crucial for the processing of syntactically complex sentences (Skeide et al., 2016; Wilson et al., 2011). The structural properties of these fiber tracts are prime candidates for the explanation of inter-individual differences in language performance and the effects of training during learning.

These properties include, but are not limited to, the trajectories and density of nerve fibers, determining which neurons may exchange information, as well as diameters and myelination of axons, impacting transmission speed as well as synchronization and ephaptic coupling between axons (Schmidt et al., 2021; Schmidt and Knösche, 2019). In living human brains, these structural properties are only indirectly accessible by imaging techniques, mainly based on magnetic resonance imaging (MRI). For example, it is believed that the T1 relaxation time directly correlates with the myelin content of the tissue (Bock et al., 2013; Stüber et al., 2014; Weiss et al., 2010). Therefore, T1 weighted as well as quantitative T1 imaging are used to chart white matter myelination. The same applies to magnetic transfer measurements (Henkelman et al., 2001). Diffusion weighted MRI protocols deliver information on the spatial trajectories of nerve fibers (see Jones et al., 2013, for a critical review), but also on other properties of the fibers, such as axonal diameter (Assaf et al., 2008; but see also Paquette et al., 2020) and myelin sheath thickness (g-ratio; see Mohammadi and Callaghan, 2021). Importantly, all these measures, and any metrics derived from them, are sensitive to a mixture of different tissue properties. Moreover, the spatial resolution of MRI (usually >1 mm) entails that the properties of millions of fibers as well as hundreds of thousands of other structures (e.g., glia cells) are averaged in each voxel (Walhovd et al., 2014).

Prior work has demonstrated white matter plasticity in multiple studies on second language learning. Many of them are cross-sectional and compare populations with and without certain second language skills (Cummine and Boliek, 2013; Hämäläinen et al., 2017; Mamiya et al., 2016; Pliatsikas and Chondrogianni, 2015; Vandermosten et al., 2015). These studies therefore target white matter plasticity occurring over a long (and not precisely defined) period of time. In contrast, Schlegel and colleagues (2012) used diffusion MRI to show reorganization of major white matter fiber tracts over a period of 9 months, during which the participants intensively learned a second language (Chinese).

Although the neural bases of first and second language acquisition are discussed to be partly overlapping (Perani and Abutalebi, 2005), there may be substantial differences, in that second language acquisition relies on more variable and widespread neural networks (Cargnelutti et al., 2019; Dehaene et al., 1997) compared to the relatively well defined first language network (Friederici, 2017, 2011). For native language acquisition, we found developmental changes in the gray and white matter of the language network (Cafiero et al., 2019; Ekerdt et al., 2020; Huber et al., 2018). In adults, the language capabilities are largely established, which is paralleled by a fully matured language network (Skeide et al., 2016). However, adults may be trained to further improve in certain aspects of their mother tongue, and the question is whether this leads to non-invasively detectable reorganization of the white matter. Learning rate related changes in language relevant gray and white matter regions were observed in adults as a function of word learning in their native language over a short period of time (<1 hour) (Hofstetter et al., 2013). Flöel and colleagues (2009) used a region-of-interest approach focused on Broca’s area and its right hemispheric homologue to identify changes in white matter as a function of artificial grammar learning. Nevertheless, it is still open whether individual differences in native language processing and learning relate to individual white matter brain structure.

Therefore, we investigate whether the behavioral differences in adults processing syntactically complex sentences in their native language are rooted in structural differences in long-range fiber connections, whether the success of intense training over a relatively short period of time is predicted by such structural traits, and whether such training would in fact induce further structural changes.

To elucidate these questions, we used an experiment, during which adult German participants were first tested on understanding thematic role assignments in complex German sentences, and then trained to improve their performance. We explored and compared structural connectivity matrices obtained by tractography from diffusion MRI data acquired before and after training and related these to behavioral performance at both time points. In the analyses, we focused on the aspect of long-range connectivity in the language network, rather than metrics that might be more sensitive to local microstructural properties, such as fractional anisotropy or mean diffusivity.

## 2. Methodology

### 2.1 Paradigm

The experiment was designed to investigate how brain function and white matter microstructure change during multi-day language training. Training was performed during four out of the five working days of one week. On the first training day, prior to the experiment, reading span (Daneman and Carpenter, 1980) and digit span (WAIS-IV, Wechsler et al., 2008) were acquired as measures for language specific and general working memory abilities, respectively. On each training day, the participants listened to 66 German center-embedded sentences: half with single and the other half with double center embedding.

#### Example for single center embedding

Ihr Freund sagte, dass Gustav, der Marlene überschätzte, Klavier spielt, um sich zu bilden. [Her friend said that Gustav, who overestimated Marlene, plays piano, in order to educate himself.]

#### Example for double center embedding

Yvonne dachte nicht, dass Bernd, der Leo, der intelligent ist, liebte, Maria verfolgen will. [Yvonne did not think that Bernd, who loved Leo, who is intelligent, wants to pursue Maria.]

Each sentence was followed by a content question probing the participant’s understanding of the thematic role assignment. The answer was recorded by delayed key press within a predefined time window and acknowledged by a visual feedback (smiley/frowny). In case of a wrong or missing answer, the same sentence was repeated and additionally displayed on screen. A different content question was then asked and again acknowledged by feedback. Irrespective of the correctness of the second answer, the experiment was continued with the next trial. During the measurements, MEG was recorded with a Neuromag Vectorview device (results reported in Wang et al., 2021).

The performance of the participants was measured as the percentage of correct answers to the first content questions (#correct / (#incorrect + #missed)).

Finally, T1, diffusion, and resting state functional MRI data were acquired three times: before the experiment (scan a), immediately after it (scan b), and 3 weeks later (scan c). See below for technical details.

### 2.2 Participants

The sample included 28 right-handed participants and inclusion criteria were as follows: subjects were 18-35 years old at the time of recruitment, they were German native speakers with normal or corrected to normal hearing and vision, and no history of substance abuse (alcohol or drugs). Subjects with neurological or psychiatric disorders, past neurosurgery, neuroactive medication, claustrophobia, pregnancy, or other contraindications for MRI were excluded. Written informed consent was obtained from all participants prior to the experiment. The study was approved by the ethics committee of the University of Leipzig.

### 2.3 MRI data acquisition

MR images were acquired with a 3T Siemens Magnetom Prisma MRI scanner. A high-resolution (1 mm^3^) structural T1-weighted scan was obtained (MP-Rage, TR=1.3 s, TE = 3.93 ms; α = 10°; 1×1×1 mm). Diffusion weighted MRI (DWI) was acquired with the standard GE-EPI protocol. The employed parameters for diffusion data were: TR = 12 s, TE = 100 ms, A/P phase encoding direction, 72 slices, FOV = 220 x 220 mm^2^, acquisition matrix 128×128, 1.7 mm isotropic voxels, 60 diffusion-weighted images (b = 1000 s/mm^2^), and 7 no diffusion weighting (b_0_) images. Multiband and fat saturation techniques were implemented to improve quality data. Finally, functional MRI was measured with gradient echo EPI (TR = 2 s, TE = 30 ms, flip angle 90°, resolution: 3×3×3 mm), during 15 minutes rest with eyes open.

### 2.4 Atlas selection and DWI pre-processing

In this study, we used the HCP-MMP1 atlas (Glasser et al., 2016), which includes 180 cortical regions per hemisphere. This freely available atlas has been created by a sophisticated machine learning approach. It combines information about cortical architecture, function, connectivity, and topography in a precisely aligned group average of 210 brains, and has been carefully cross-validated. It is therefore arguably one of the most comprehensive and reliable human brain atlases available today. After executing an intra-subject cross-modal registration, based on a rigid body transformation, we performed a projection of this atlas onto each participant’s MNI-T1 image. Therefore, the outcome for each subject was the HCP-MMP1 atlas co-registered with diffusion data. In addition, in order to avoid spurious inter-hemispheric connections, we added to the analysis a 5-region-parcellation of the corpus callosum from a white matter parcellation. This step was executed using the Freesurfer software.

After visual inspection for large artifacts, diffusion data were corrected from susceptibility-induced distortion, subject motion, and artifacts due to eddy currents. To estimate diffusion parameters at each voxel, Bayesian inference was performed through the tool *bedpostx*, which also resolves voxels with crossing fibers. Subsequently, probabilistic tractography was applied to reconstruct sample streamlines using the *probtrackx2* command with default parameters (5000 samples per seed voxel, maximum of 2000 steps per streamline, curvature threshold of 0.2, step length of 0.5 mm). In order to identify experimental effects related to language training, tractography was performed separately for each hemisphere. This was done in order to avoid potential misidentifications of connections when tracking through the highly convergent corpus callosum region. This way, we generated a seed-to-seed connectivity matrix per subject and hemisphere. The matrix entries *C_ij_* correspond to the number of streamlines generated from region *i* (source) and entering region *j* (target). Both tools, *bedpostx* and *probtrackx2*, are part of the FSL software.

### 2.5 Analysis of Connectivity

Based on the connectivity matrices described above, and the two issues raised in the introduction, we investigated the following relations:

The first issue concerns the relation between the brain structural precondition and performance prior to training.

(Q1) Does the *a priori* connectivity (scan a, before training) correlate with the initial performance (day 1)?
(Q2) Does this *a priori* connectivity (scan a) correlate with the performance change over the 4-days-experiment (day 4 minus day 1, training effect)?

The second main question concerns the relation between brain structural changes and training induced performance change with two sub-questions:

(Q3) Does the change in connectivity over the experiment (the difference in connectivity values between scan b minus scan a) correlate with the performance change over the 4-days-experiment (training effect, day 4 - day 1)?
(Q4) Although we focus on individual differences, we will nonetheless ask whether there is a group-level change in connectivity between the time points before and after the 4-days of training (scans b versus a).

Because the behavioral training effect was most prominent for the double center embedded sentences (see Results), we used those trials in the connectivity analysis. For each of the above questions, we performed statistical tests on three different types of structural connectivity data:

i. *Connection based analysis* performed for each entry of the connectivity matrix.
ii. *Node based analysis* performed on each node of the network, calculating a centrality metric that adds up streamlines generated from a source region, and another which quantifies streamlines that reach a target region.
iii. *Network based analysis* performed on the relative prevalence of network motifs obtained by singular value decomposition (see below for details).

Since the connectivity values cannot be considered normally distributed, we exclusively used non-parametric tests. Specifically, we applied Spearman’s correlation, corrected for multiple comparisons by the false discovery rate criterion proposed by Benjamini and Hochberg (FDR-BH) (Benjamini and Hochberg, 1995). For the pairwise comparison in Q4, we performed Mann-Whitney U-tests, also followed by FDR-BH correction.

The network based analysis (iii) is motivated by the notion that in the brain network nodes and connections do not act in isolation, but as part of larger (sub-)networks. Consequently, any kind of specialization, either occurring during a live-time development (relevant for Q1, Q2) or induced by an intense training process (relevant for Q3, Q4) should also involve coherent changes of entire network patterns, or motifs. Therefore, we applied singular value decomposition (SVD) to factorize the whole structural connectome of each subject and hemisphere, and identify any network component associated with the language training process. Separately for the two hemispheres, the structural matrices of all subjects and scans were arranged as column vectors, forming a matrix *M* which was decomposed into left singular vectors *U*, singular values *Σ*, and right singular vectors *V*: *M* = *UΣV^T^*. The left singular vectors represent orthogonal and normalized network motifs, the singular values indicate their prevalence across all structural matrices, and the right singular vectors indicate the relative prevalence of each network motif in each subject and scan. Each column of the *V* matrix can now be used to perform the statistical tests and correlation analyses described above.

Connectivity matrices, and relevant Matlab’s code used for SVD analysis and plotting are available on a GitHub repository at https://github.com/hschmidt82/Yerevan_public.

## 3. Results

### 3.1 Behavioral

Figure 1 displays the mean performances over subjects for each training day, separately for single and double embedding sentences. These results, which were already reported and discussed elsewhere (Wang et al., 2021), show a significant performance improvement for both types of sentences, but for the simpler sentences this improvement had to be small since the initial performance was already quite high. Due to this ceiling effect, we decided to use for the connectivity analyses the double embedded sentences only.

**Figure 1:**
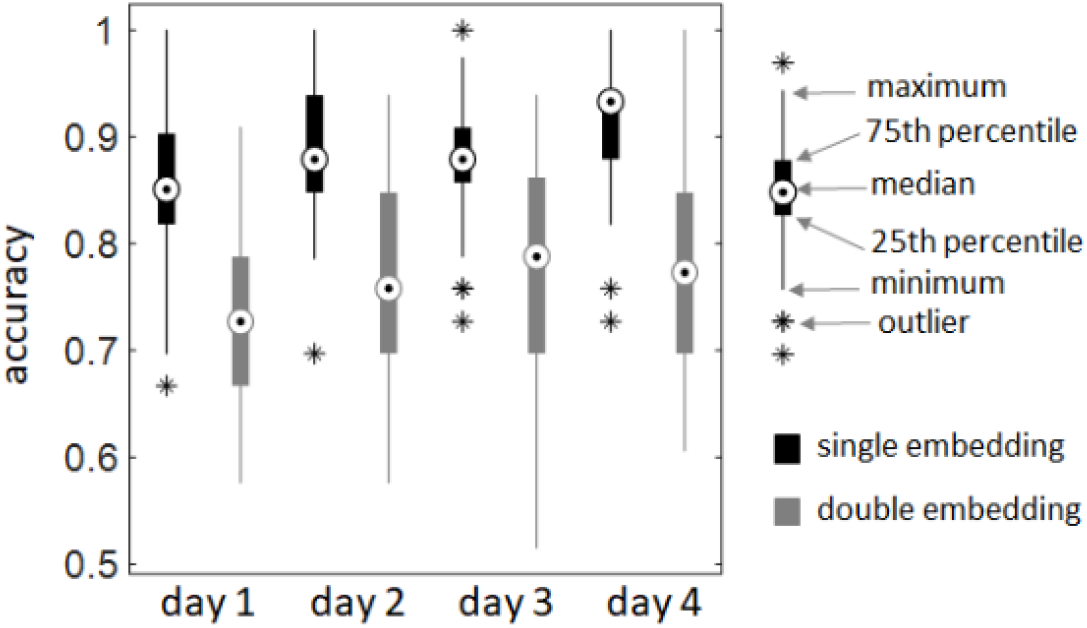
Behavioral data. For both, single and double embedding, there is a significant difference between day 4 and day 1 (p < 0.005, FDR corrected). For the double embeddings, there are also significant differences between day 1 and day 2 as well as between day 1 and day 3 (p < 0.05, FDR corrected).

In order to probe how the initial performance was related to the participants’ memory abilities, we correlated it with reading span and digit span (levels reached, forward and backward averaged). Reading span showed a significant correlation with the performance on single (*r* = 0.31, *p* < 0.05), but not on double embedded sentences (where performance was initially quite low). Digit span did not yield any significant correlation.

### 3.2 Left Hemisphere

Both the connection based and the node-based analyses did not yield any statistically significant correlations between either of these two approaches and behavioral data (Q1-Q3). In addition, we did not find any group level differences between before and after training (Q4).

In contrast, the network based analysis yielded two relevant network motifs related to behavioral performance. The first motif appears to exhibit a predisposition effect, as its prevalence in the subjects correlates with their performance on the first day (significant for scans a and b, marginally significant with *p* < 0.075 in scan c; see Figure 2). This gives a direct answer to question Q1. Figure 2 displays the main areas and connections of this network component. It comprises areas in the medial prefrontal cortex (10v, 9m, d32), the posterior cingulate cortex (23c, DVT [dorsal visual transition area], POS2 [parietal occipital sulcus]), the frontal operculum (FOP2), and the temporo-parieto-occipital junction (PSL [perisylvian language area], STV [superior temporal visual area]). While some of these areas are known for their specific involvement in language processing (particularly PSL), others have been reported to be activated in theory of mind (watching socially interacting objects; areas 10v, 9m, PSL, STV), motor functions (9m, 23c, FOP2, POS2, PSL, STV, DVT), and working memory tasks (9m [faces], d32 [all images], 23c [body], DVT [places], POS2 [body, faces]) (Glasser et al., 2016).

**Figure 2:**
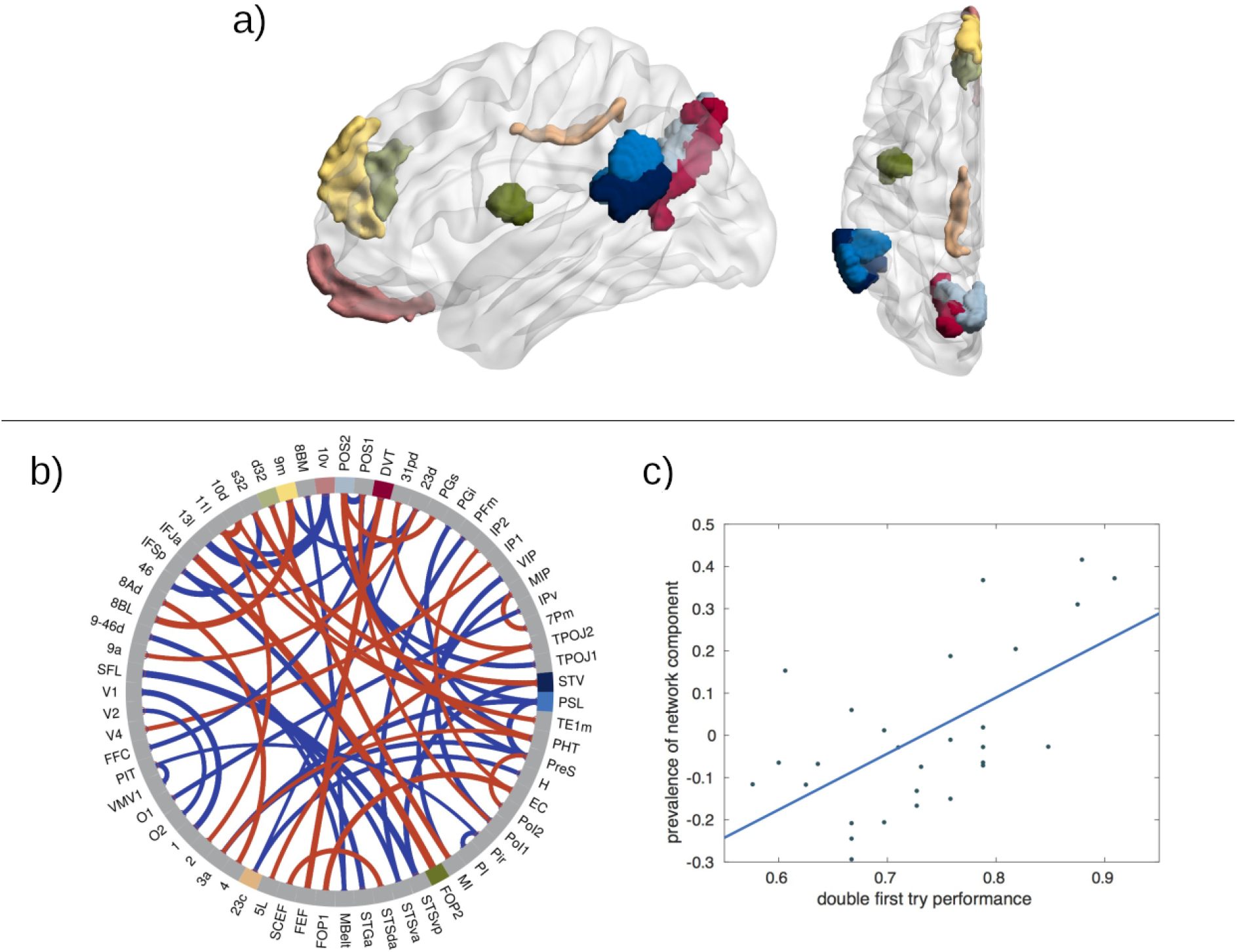
Network motif 1 in the left hemisphere, whose prevalence before the training correlates with the baseline performance on day 1. (a) Sagittal and axial view (created using BrainNet Viewer; Xia et al., 2013) of nine brain areas where the network component has highest (absolute) connectivity (labels according to Glasser et al., 2016, in panel b). (b) Chord plot of strongest connections in the network motif (red: positive, blue: negative), with line thickness indicating strength of connection. Note that connections that have negative weightings in the network motif actually correlate negatively with the motif prevalence. 67 of the 185 brain areas are plotted, the main constituents of the network motif (panel a) are highlighted in color. (c) Regression plot of network prevalence (right singular vector) against double first try performance (r = 0.567, p = 0.046).

The second motif indicates structural changes related to the training process. Its difference in prevalence between scan b (after training) and scan a (before training) significantly correlates with the change in performance between the last (day 4) and the first (day 1) days of training (Figure 3). This finding directly relates to question Q3. Figure 3 shows the main areas and connections of this network component. They partially overlap with the first network motif (areas 9m, 23c, POS2, STV, and PSL). In addition, this network motif includes parts of auditory association cortex (TA2 on planum polare, STSvp in superior temporal sulcus), dorsolateral prefrontal cortex (9p), hippocampus (H), and ventral stream visual cortex (PIT [posterior inferior temporal]). Again, these areas have been reported in a wide variety of experimental conditions, including language processing (PSL, TA2, STSvp, H), theory of mind (watching socially interacting objects; areas 9m, 9p, PSL, STV, STSvp), motor functions (9m, 23c, POS2, PSL, STV, STSvp), and working memory tasks (9m [faces], 23c [body], POS2 [body, faces], H [all], PIT [faces]), (Glasser et al., 2016).

**Figure 3:**
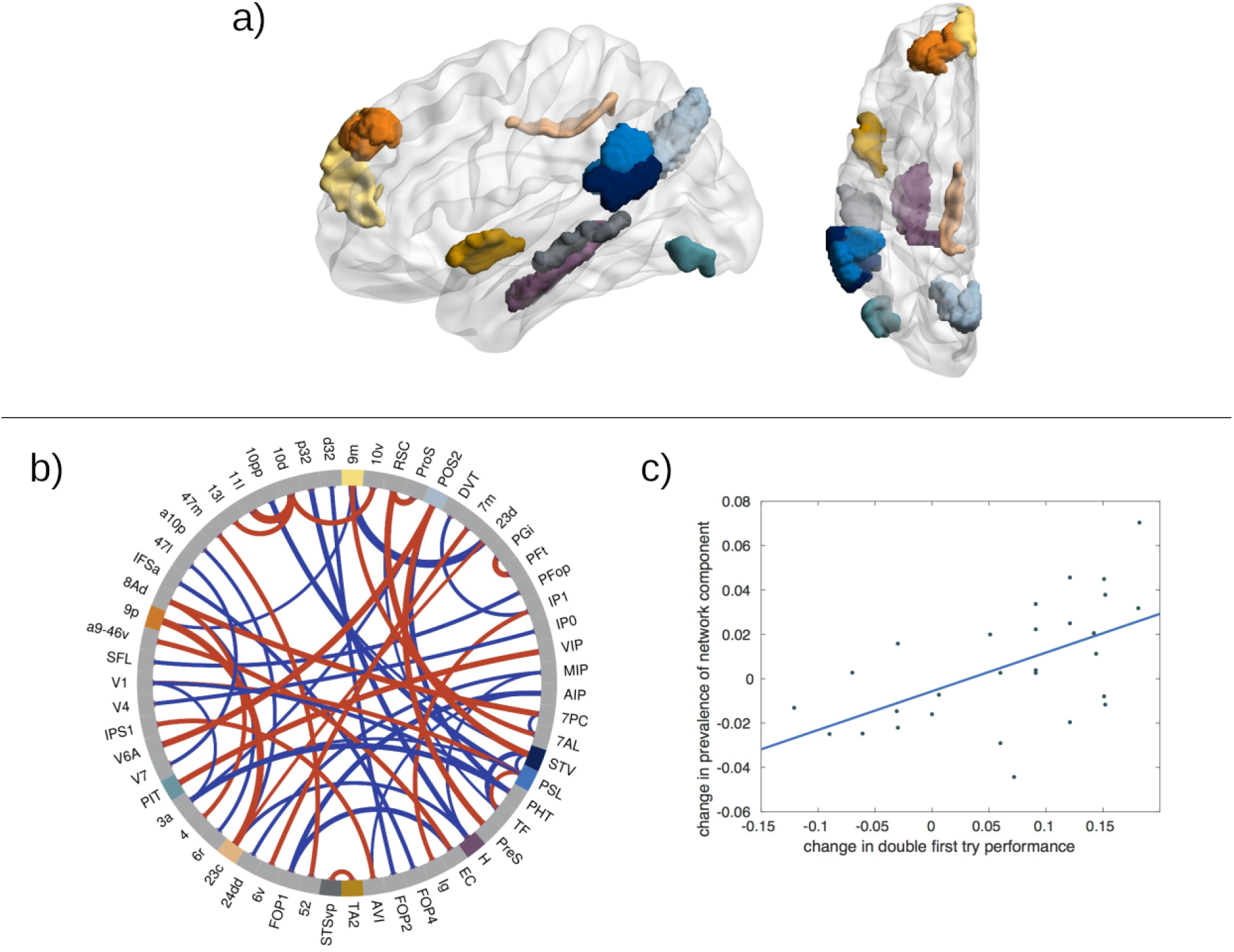
Network motif 2 in the left hemisphere, whose changes over the training period correlate with the performance change between day 1 and day 4. (a) Sagittal and axial view (created using BrainNet Viewer; Xia et al., 2013) of ten brain areas where the network component has highest (absolute) connectivity (labels according to Glasser et al., 2016, in panel b). (b) Chord plot of strongest connections in the network motif (red: positive, blue: negative), with line thickness indicating strength of connection. Note that connections that have negative weightings in the network motif actually correlate negatively with the motif prevalence. 59 of the 185 brain areas are plotted, the main constituents of the network motif (panel a) are highlighted in color. (c) Regression plot of change in network prevalence (right singular vector) against change in double first try performance (r = 0.606, p = 0.035).

None of the two left hemispheric motifs yielded significant results concerning question Q2 (prediction of training effect by pre-experimental connectivity) and Q4 (group level change of connectivity through training).

### 3.3 Right Hemisphere

Also in the right hemisphere, connection based and the node-based analyses did not result in any statistically significant differences or correlations.

The network-based analysis also did not yield any structural changes related to the training process (Q2-4). However, a motif was identified whose prevalence correlates with the subjects’ performance on the first day, thus relating to question Q1. Figure 4 displays the main areas and connections of this network component. This network motif appears very different from the left-hemisphere motifs 1 and 2, and mainly comprises inferior parietal cortex (PGs, PGi, PFm), the temporo-parieto-occipital junction (TPOJ1), primary auditory cortex (RI [retro-insular]), premotor cortex (6r), and visual cortex (V1, V3, V3A, PH). While for some of the inferior parietal areas, a specific involvement in language processing has been reported (PGi, TPOJ1), others are known to be specifically deactivated for language (PGs, PFm) (Glasser et al., 2016). Interestingly, the most strongly represented part of the primary auditory cortex, area RI, is reported as deactivated during language processing, while the other core and belt areas are strongly activated in the same task (Glasser et al., 2016, Fig. 12).

**Figure 4:**
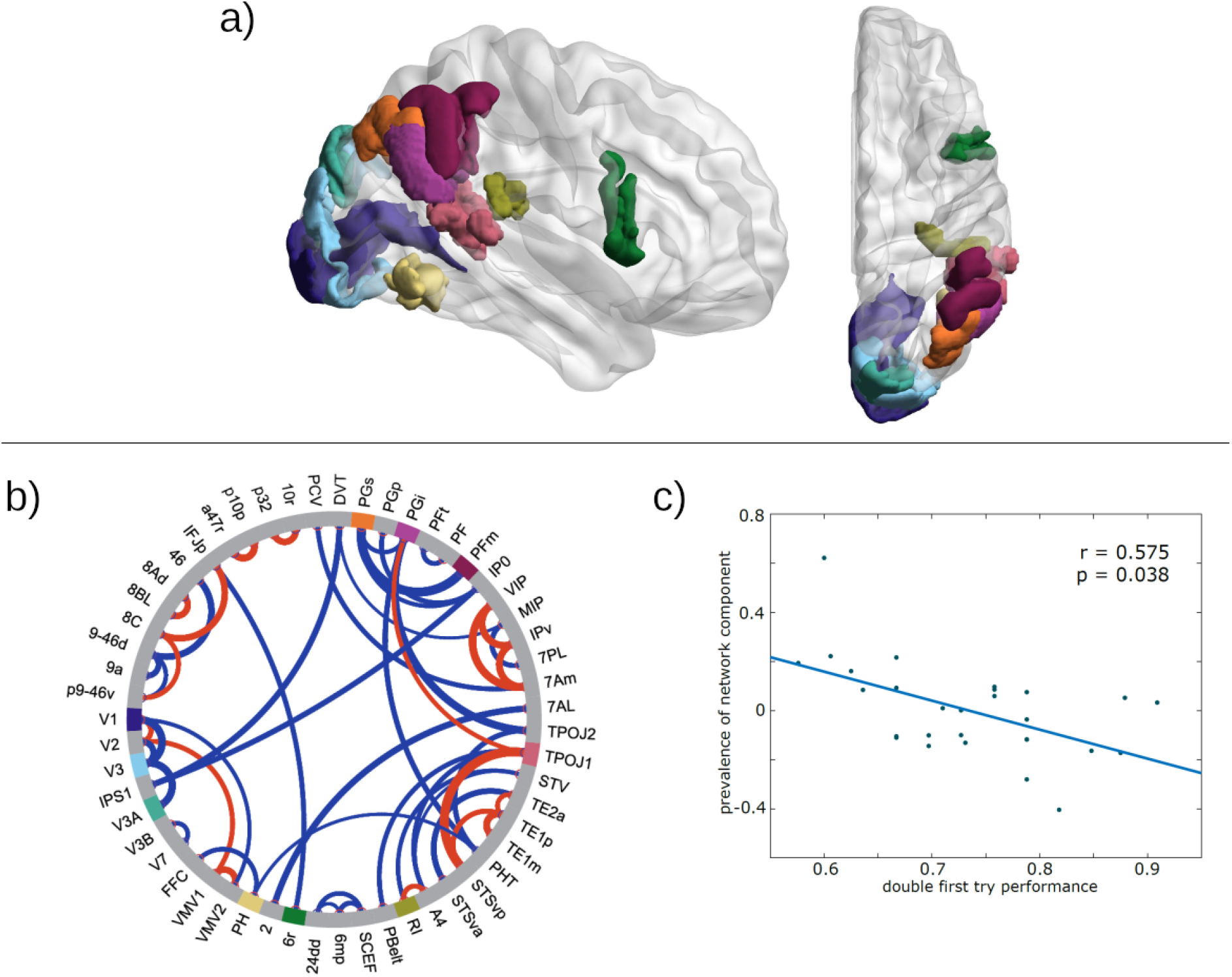
Network motif 3 in the right hemisphere, whose prevalence before the training correlates with the baseline performance on day 1. (a) Sagittal and axial view (created using BrainNet Viewer; Xia et al., 2013) of nine brain areas where the network component has highest (absolute) connectivity (labels according to Glasser et al., 2016, in panel b). (b) Chord plot of strongest connections in the network motif (red: positive, blue: negative), with line thickness indicating strength of connection. Note that connections that have negative weightings in the network motif actually correlate negatively with the motif prevalence. 55 of the 185 brain areas are plotted, the main constituents of the network motif (panel a) are highlighted in color. (c) Regression plot of network prevalence (right singular vector) against double first try performance (r = 0.575, p = 0.038).

## 4. Discussion

In this study, our goal was to elucidate the relationship between individual white matter brain anatomy, as revealed by diffusion tractography, and the comprehension of syntactically complex sentences in adult native speakers. We were asking if and how the given structure of the white matter of an individual influences their ability to extract thematic roles from center embedded sentences (Q1), and how it affects the improvement of that ability through intense training (Q2). Moreover, we also sought to answer the question whether intense training can induce measurable white matter changes. Specifically, we tested if such changes are related to the individual performance improvement (Q3), and if there was a general (group-level) difference in white matter connectivity between the time points before and after training (Q4). Each of these questions was approached based on connections between nodes (connection based), the degree of nodes (node based), and the prevalence of network motifs (network based).

First of all, none of the tested correlations and differences did render significant at the connection or node levels, where massive correction dictated by the high number of tests might have obscured more subtle effects. It seems reasonable to assume that this type of testing leads to overcorrection, since it is unlikely that each of the many network connections varies independently. Instead, entire subnetworks might react as a whole. This notion motivated our network analysis, and indeed we were able to identify three network motifs, two of which (one in each hemisphere) predicted the initial language performance of the participants, while the third motif (in the left hemisphere) changed during training in correlation with the individual performance change.

We will now discuss these data in more detail. The first main question concerns the relevance of the individual pre-existing white matter connectivity for language performance and training success in the native language. Here, we can conclude: yes, there are global structural properties of the white matter connectome, which predict the individual performance in comprehending complex sentences, and these properties can be extracted from diffusion MRI by tractography (Q1). In the left hemisphere, the identified network motif (Figure 2) is widely spread over the cortex and involves areas that are known to be engaged in a variety of different brain functions. It contains connections that correlate positively or negatively with the performance, respectively. Positive correlations were mainly found for areas known for their involvement in working memory tasks (9m, d32, DVT, POS2), while some areas related to language and theory of mind tended to correlate negatively with the individual performance (PSL, 10v). This suggests that the different initial performances of the participants (67…100 % for single embedded sentences and 58…91 % for double embedded sentences) are rooted in differences in their working memory system. This presumption is further supported by the finding that the initial performance on single embedded sentences, which was already quite high prior to training in most subjects, correlated with the individual reading span score. The fact that reading span, but not digit span, showed such correlation might hint that language specific rather than general working memory is relevant here. These interpretations are interesting given that psycholinguistic theory has proposed that the ability to deal with syntactically complex sentences is influenced by the individual working memory capacity (Just and Carpenter, 1992; MacDonald et al., 1992).

In addition, there was also a right hemispheric network motif, the prevalence of which predicted the individual performance before the training (Figure 4). In contrast to the motif in the left hemisphere, the right hemispheric motif had a strong focus on modality-specific cortices (visual, auditory, motor), and additionally involved many working memory specific areas (PGs, PFm, 6r).

These very same network motifs (or any other network motif), however, did not predict the ability of the participants to improve their performance through training (Q2). Naturally, it must remain open whether this is because the individual training abilities are governed by other factors than those reflected by white matter tractography, or because the statistical power for the correlation with training induced performance improvement was not sufficient.

Remarkably, the network motifs found in this study did not exhibit any prominent contribution from the syntax-related Broca’s area (areas 44 and 45 in the Glasser atlas). White matter connections between area 44 and the posterior temporal cortex were shown to correlate with performance on processing complex sentences during development (Skeide et al., 2016). In adults, the white matter connection with that area has been found to correlate with individual performance in an artificial grammar-learning task using a region-of-interest approach (Flöel et al., 2009). Here, we find that individual performance differences of adults processing syntactically complex sentences of their native language mainly rest on differences in working memory-related regions rather than syntax-related regions.

The second main question investigated here focused on training and thereby learning related changes in white matter properties. We could show that the individual performance change over the training period significantly correlated with the individual prevalence change of a particular network motif in the left hemisphere (Figure 3). This finding suggests that individual learning success for processing complex sentences depends on some reorganization of the white matter within the left hemisphere (Q3). At the group level, our analyses did not reveal significant structural changes in the white matter (Q4), although the participants significantly improved their performance at the group level (Figure 1). In other words, averaging across the individuals’ white matter structure could not explain the observed group performance difference; rather it was the individual brain structure, which provided an explanation.

Interestingly, the motif related to individual performance change bears substantial similarity to the one that predicts the initial performance (see above), but also features some differences. The most striking difference is the additional involvement of temporal areas (TA2, STSvp; compare Figures 2a and 3a). Most of the strong connections show positive correlations with the training success, involving areas reported for language (TA2, STSvp), theory of mind (9m, 9p, STV, STSvp), working memory (9m, 23c, POS2, PIT) and motor functions (9m, 23c, POS2, STV, STSvp). Prominent areas with negative connections were PSL (involved in various functions, including language) and H (hippocampus). This picture seems to suggest that the training process is associated with some reorganization of widespread networks involved in multiple functional aspects of the brain.

## 5. Conclusion

In summary, we can state that the individual modulation of the ability to extract thematic roles from center-embedded sentences seems to depend on widespread networks of white matter fibers connecting areas in different parts of the cortex. The initial individual level of language performance, which has been acquired throughout life, seems to depend mainly on (temporo-)parietal, medial prefrontal, frontal opercular, and cingulate areas, many of which are known for their involvement in working memory abilities. The rapid performance improvement of processing syntactically complex sentences induced by the relatively short, but intense, training during the experiment appears to induce changes in a diverse network additionally involving medial and superior temporal regions, spanning language, working memory, theory of mind, and motor functions. Taken together, this suggests that under the pressure of the training, subjects used a multitude of strategies to improve their performance, but it appears that in the long run, working memory is the key ability to master complex center-embedded sentences.

The present data suggest that individual language performance can be explained by individual white matter structural patterns – a relation, which may hold for individual differences observed in cognitive functions more generally. Thus, the individual differences in the brain’s white matter structure may be a crucial factor to be considered when discussing variations in cognitive performance.

## Acknowledgments

We thank the Bunge y Born Foundation – Williams Foundation – Max Planck Institute for supporting SMS with a scholarship. At the time of this study, SMS was a doctoral fellow at CONICET Argentina. HS was supported by a grant of the German Research Foundation awarded to TRK (KN 588/7-1).

